# Widespread use of the “ascidian” mitochondrial genetic code in tunicates

**DOI:** 10.1101/865725

**Authors:** Julien Pichon, Nicholas M. Luscombe, Charles Plessy

**Affiliations:** Okinawa Institute of Science and Technology Graduate University 1919-1 Tancha, Onna-son, Kunigami-gun Okinawa, 904-0495, Japan; Université de Paris; The Francis Crick Institute, 1 Midland Road, London, NW1 1AT, UK; UCL Genetics Institute, University College London, Gower Street, London, WC1E 6BT, UK

**Keywords:** Tunicate, Oikopleura, Genetic code, Mitochondria, Cytochrome oxidase sub-unit I

## Abstract

**Background:** Ascidians, a tunicate class, use a mitochondrial genetic code that is distinct from vertebrates and other invertebrates. Though it has been used to translate the coding sequences from other tunicate species on a case-by-case basis, it is has not been investigated whether this can be done systematically. This is an important because a) some tunicate mitochondrial sequences are currently translated with the invertebrate code by repositories such as NCBI’s GenBank, and b) uncertainties about the genetic code to use can complicate or introduce errors in phylogenetic studies based on translated mitochondrial protein sequences.

**Methods:** We collected publicly available nucleotide sequences for non-ascidian tunicates including appendicularians such as *Oikopleura dioica*, translated them using the ascidian mitochondrial code, and built multiple sequence alignments covering all tunicate classes.

**Results:** All tunicates studied here appear to translate AGR codons to glycine instead of serine (invertebrates) or as a stop codon (vertebrates), as initially described in ascidians. Among Oikopleuridae, we suggest further possible changes in the use of the ATA (Ile → Met) and TGA (Trp → Arg) codons.

**Conclusions:** We recommend using the ascidian mitochondrial code in automatic translation pipelines of mitochondrial sequences for all tunicates. Further investigation is required for additional species-specific differences.

## Introduction

Tunicates are marine animals that have acquired the capacity to produce cellulose by horizontal gene transfer approximately 500 million years ago (Matthysse et al., 2004; Nakashima et al., 2004). Together with vertebrates and cephalochordates, they belong to the chordate phylum, in which they share morphological features such as a muscular tail during larval stages. Phylogenetic studies place the tunicates as the closest living relatives of vertebrates (Delsuc et al., 2006). Tunicates can be subdivided in three classes: Thaliacea (free-swimming colonial species, for instance salps or dolioids), Appendicularia (free-swimming solitary species with an adult morphologically similar to the larval stage of other tunicates), and Ascidiacea (attached to solid substrates in their adult stage, for instance sea squirts). The relationship between these classes and therefore their mono- or paraphyly has been revised multiple times. For instance the 18S rRNA analysis of Stach and Turbeville (2002) nested Appendicularia within Ascidiacea, but more recently Delsuc et al. (2018) placed them as sister groups using a multigene approach. The paraphyly of Ascidiacea is now widely accepted, as the above studies and others demonstrated that they contain the Thaliacea.

Mitochondrial genomes undergo major changes at the geological time scale due to their small size and clonal reproduction, including changes to their genetic code (Osawa et al., 1992). The first evidence that ascidians use a specific mitochondrial genetic code stemmed from observations that the Cox1 sequence from *Halocynthia roretzi* (Yokobori et al., 1993) and the Cox3 sequence of *Pyura stolonifera* (Durrheim et al., 1993) are interrupted by stop codons if translated using the vertebrate mitochondrial code. Reassignment of AGR codons to glycine was later confirmed by the discovery of a glycine (Gly) tRNA in the *H. roretzi* genome (Yokobori et al., 1999) and by the sequencing of its anticodon (U*CU) (Kondow et al., 1999). Apart from the AGR codons, the ascidian code is similar to the vertebrate and the invertebrate ones, with ATA assigned to methionine (Met) and TGA to tryptophan (Trp) (Yokobori et al., 1993).

This genetic code is known as the “ascidian” genetic code; however, it is also used by non-ascidian tunicates, such as the thaliacean *Doliolum nationalis* (Yokobori et al., 2005). The possibility that this genetic code emerged earlier than tunicates was raised by the study of partial genome sequences of *Branchiostoma lanceolatum* (Delarbre et al., 1997) leading to the proposition that AGR might encode Gly in cephalochordates. While this seemed to be supported by the discovery of a putative TCT (Gly) tRNA in the full mitochondrial genome of *B. lanceolatum* (Spruyt et al., 1998), this hypothesis was later ruled out by an analysis of the related amphioxus *Branchiostoma floridae* (Boore et al., 1999), and has not been reconsidered since. Finally, studies on the appendicularian branch showed compatibility between the mitochondrial sequence of *Oikopleura dioica* and the ascidian code (Denoeud et al., 2010). Nevertheless, support for compatibility was not demonstrated explicitly for the ATA and TGA codons and the mitochondrial sequence of *O. dioica* were not released in International Nucleotide Sequence Database Collaborations (INSDC) databanks.

The cytochrome c oxidase subunit 1 (Cox1) is the most conserved mitochondrial protein. Although no mitochondrial genome has been fully sequenced yet for appendicularians, partial Cox1 sequences are present in the INSDC databanks for Oikopleuridae. Sakaguchi et al. (2017) reported that all *Oikopleura dioica* mitochondrial sequences (AY116609–AY116611 and KF977307) may be contaminations from bacteria or cnidarians, and provided partial sequences for *Oikopleura longicauda* in the same study. Partial mitochondrial sequences were published for *Bathochordaeus* and *Meso-chordaeus* species by Sherlock et al. (2017). In addition, Naville et al. (2019) recently published draft genome for several appendicularian species. Therefore, to assess whether the ascidian mitochondrial code is used across the whole tunicate subphylum, we took advantage of these public data and prepared a curated alignment of Cox1 sequences comprising representatives of the major tunicate branches, to study the consensus sequences at conserved residues.

## Methods

We identified Cox1 and Cytochrome b (Cob) gene sequences for *Oikopleura longicauda*, *Mesochordaeus erythrocephalus* and *Bathochordaeus stygius* by screening published genome assemblies (Naville et al., 2019) with the partial Cox1 sequence of *O. longicauda* `LC222754.1` (Sakaguchi et al., 2017) using `tblastn` and the ascidian mitochondrial code (`-db_gencode=13`) (Gertz et al., 2006). Mitochondrial genome sequences were then translated using the `cons` and `getorf` commands from EMBOSS (Rice et al., 2000), using the ascidian mitochondrial code.

### Oikopleura longicauda

We identified the circular contig `SCLD01101138.1` (length: 10,324 nt) as a potential mitochondrial genome, and translated Cox1 from position 4530 to 6230. We also translated Cob from 3697 to 4668.

### Mesochordaeus erythrocephalus

We translated Cox1 in contig `SCLF01725989.1` (length 7,034 nt) on reverse strand from position 1792 to 272. Using the same method with *O. longicauda*’s Cob sequence as a bait, we also recovered a Cob sequence from contig `SCLF01109548.1` (length 5,010 nt), reverse strand, 1604 to 2590.

### Bathochordaeus stygius

We used the consensus of the published *B. stygius* Cox1 sequences `KX599267.1` to `KX599281.1` from GenBank (Sherlock et al., 2017), to screen the genome and scaffold `SCLE01415711.1` (length 10,388 nt) gave a perfect hit. We translated Cox1 from position 8054 to 6522 on the reverse strand, and a partial Cob sequence from scaffold `SCLE01415711.1` (2319 to 2963, reverse strand). We also found a second fragment aligning well with C-terminal sequences between positions 2373 and 1978, but we did not include it due to the difficulty of resolving the overlap between both fragments. When screening with the *M. erythrocephalus* Cox1 sequence recovered above, we found that another scaffold `SCLE01416475.1` gave a perfect hit, hinting at a possible contamination.

### Oikopleura dioica

To assemble a Cox1 sequence in *O. dioica*, we downloaded ESTs (file `10_ESTall.txt`) from Oikobase (Danks et al., 2012) and extracted hits matching the *O. long-icauda* sequence using `tblastn` (see above). We then aligned and visualised the hits using Clustal Omega (Sievers et al., 2011) and SeaView (Gouy et al., 2009), filtering out those too short or introducing gap columns. Inspection of the alignment let us notice three possible haplotypes. We generated a consensus for each of them, translated them (see above) and trimmed the proteins sequences in order to match the length of the other reference sequences in the alignment. All variants found between the haplotypes were synonymous codons. We used the same methodology to generate a consensus for Cob and translate it.

## Cox1 accession numbers

*Bathochordaeus charon* KT881544.1 ORF2 translated with ascidian code; *Bathochordaeus stygius*: SCLE01415711.1[8054:6522] translated with ascidian code; *Branchiostoma lanceolatum*: BAD93656.1; *Caenorhabditis elegans*: NP 006961.1; *Ciona intestinalis*: CAL23359.2; *Clavelina oblonga*: YP 009029840.1; *Doliolum nationalis*: BAD86512.1; *Halocynthia roretzi*: NP 038239.1; *Mesochordaeus erythrocephalus*: SCLF01725989.1[1915:260] translated with ascidian code; *Mus musculus*: NP 904330.1; *Oikopleura dioica*: consensus of Oikobase contigs KT0AAA24YA11, KT0AAA22YO17, KT0AAA22YO04, KT0AAA13YK14, KT0AAA18YK22, KT0AAA16YP04, KT0AAA13YE23, KT0AAA8YH10, KT0AAA4YK01, KT0AAA24YE23, KT0AAA18YO18, KT0AAA3YP19, KT0AAA10YF12; *O. longicauda*: SCLD01101138.1[4678:6230] translated with ascidian code; *Salpa thompsoni*: BBB04277.1.

## Cob accession numbers

*Bathochordaeus stygius*: SCLE01415711.1[2963:2319] translated with ascidian code; *Branchiostoma lanceolatum*: BAD93666.1; *Caenorhabditis elegans*: NP 006958.1; *Ciona intestinalis*: CAL23352.2; *Clavelina oblonga*: YP 009029843.1; *Doliolum nationalis*: BAD86520.1; *Halocynthia roretzi*: NP 038246.1; *Mesochordaeus erythrocephalus*: SCLF01109548.1[1604:2590] translated with ascidian code; *Mus musculus*: NP 904340.1; *Oikopleura dioica*: consensus of Oikobase contigs KT0AAA23YJ17, KT0AAA16YJ22, KT0AAA17YO14, KT0AAA10YI15, KT0AAA18YI18, KT0AAA11YF07, KT0AAA10YG05, KT0AAA1YH02, KT0AAA12YH10, KT0AAA12YC07, KT0AAA12YC07, KT0AAA18YM15; *O. longicauda*: SCLD01101138.1[3697:4668] translated with ascidian code; *Salpa thompsoni*: BBB04269.1.

## Sequence alignments

Translated Cox1 and Cob sequences were aligned using Clustal Omega (Sievers et al., 2011) and SeaView (Gouy et al., 2009). The alignments were postprocessed using the `showalign -show=n` command of EMBOSS (Rice et al., 2000) to show the differences to the inferred consensus. Graphical processing of the alignments were performed with Jalview (Waterhouse et al., 2009). The codon sequences encoding for Cox1 and Cob of the tunicate species were then added aligned to the corresponding amino-acid (three lines per species, see supplemental material) and then the text files were transposed, so that each line would correspond to a single position in the alignment, and interrogated with custom Unix commands to compute the tables presented in this manuscript.

## Results

### AGR encodes for Gly across all tunicates

We selected species according to sequence availability and to ensure coverage of the tunicate subphylum in a way that stays broad under the various hypotheses of monophyly or paraphyly for its major groups. For ascidians, we have included the phlebobranchian *Ciona intestinalis*, the aplousobranchian *Clavelina oblonga* and the pyurid stolidobranchian *Halocynthia roretzi*. For thaliaceans, we selected *Doliolum nationalis* and *Salpa thompsoni*. For appendicularians we selected *Oikopleura dioica*, *Oikopleura longicauda*, *Bathochordaeus stygius* and *Mesochordaeus erythrocephalus*. We ensured that all tunicate sequences were translated with the ascidian mitochondrial genetic code. Lastly, we included outgroup sequences from *Caenorhabditis elegans* and *Branchiostoma lanceolatum* (invertebrate mitochondrial code) and from *Mus musculus* (vertebrate mitochondrial code) to better highlight conserved amino acid positions. In Figure 1, we illustrate the relation between these species based on the phylogeny of Naville et al. (2019) for appendicularians and of Delsuc et al. (2018) for the other tunicates. We prepared Cox1 sequences from the selected species using mitochondrial genomes (for ascidians, thaliaceans, and outgroups), from draft genomes in which we found a putative mitochondrial contig after screening with a partial or a related Cox 1 sequence (for *O. longicauda*, *B. stygius*, and *M. erythrocephalus*) and from EST sequences (for *O. dioica*). We aligned the translated Cox1 and Cob sequences (Figure 2) and inspected the positions where all species use the same amino acid. Conserved glycines supported the use of AGR codons across the whole tunicate clade. We confirmed this observation with Cytochrome b (Cob) sequences obtained with the same method.

**Figure 1:**
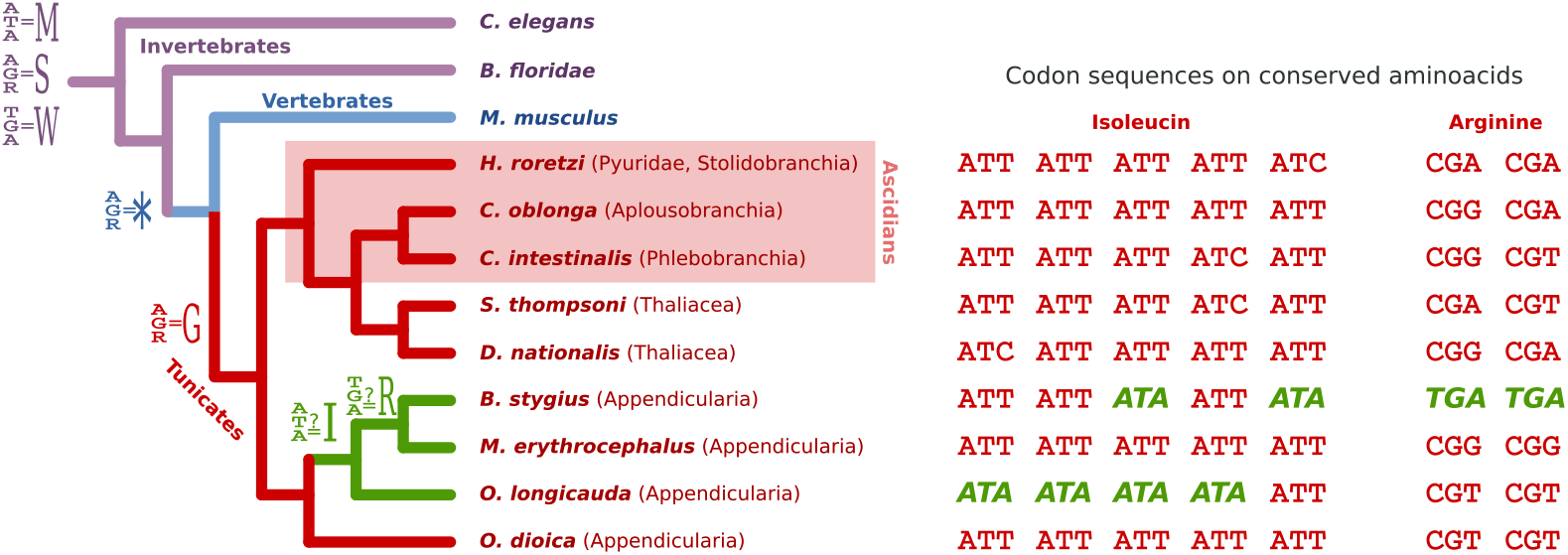
Left: Cladogram illustrating the relations between the species selected in study. Different branch colors indicate different mitochondrial genetic codes. Codon assignments with an equal sign indicate how the nucleotide sequences were translated. Codon assignments with a question mark indicate a possible finding, but were not used for translation. Ascidians, in which the AGR to Gly codon reassignment was initially discovered, are highlighted among the tunicates. Right: codon sequence of Cox1 genes on positions where proposed changes of genetic code would make all species use the same amino acid.

**Figure 2:**
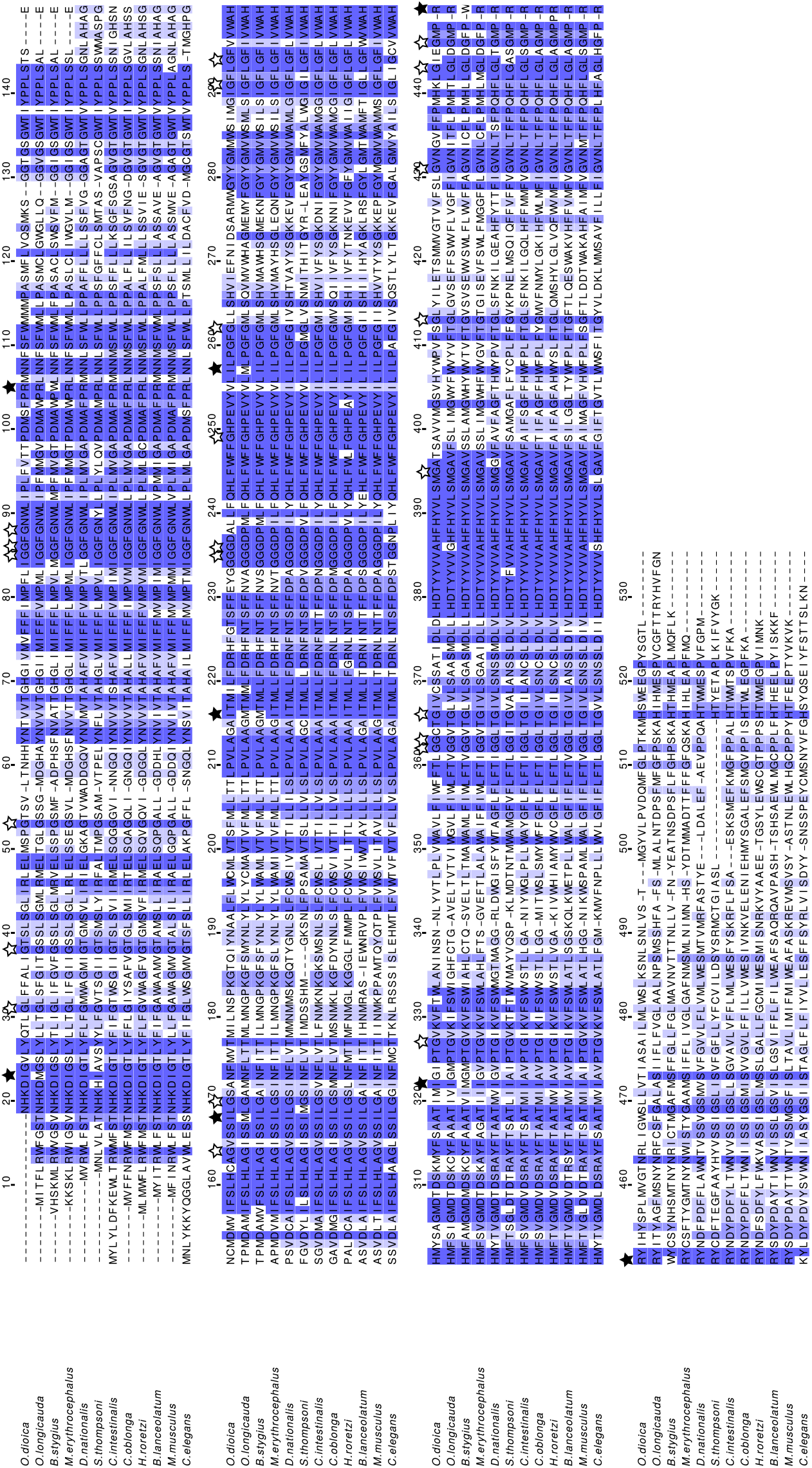
Sequence alignment of Cox1 proteins. White stars indicate conserved cysteines when at least one tunicate uses an AGR codon. Black stars indicate positions suggesting a different genetic code.

**Figure 3:**
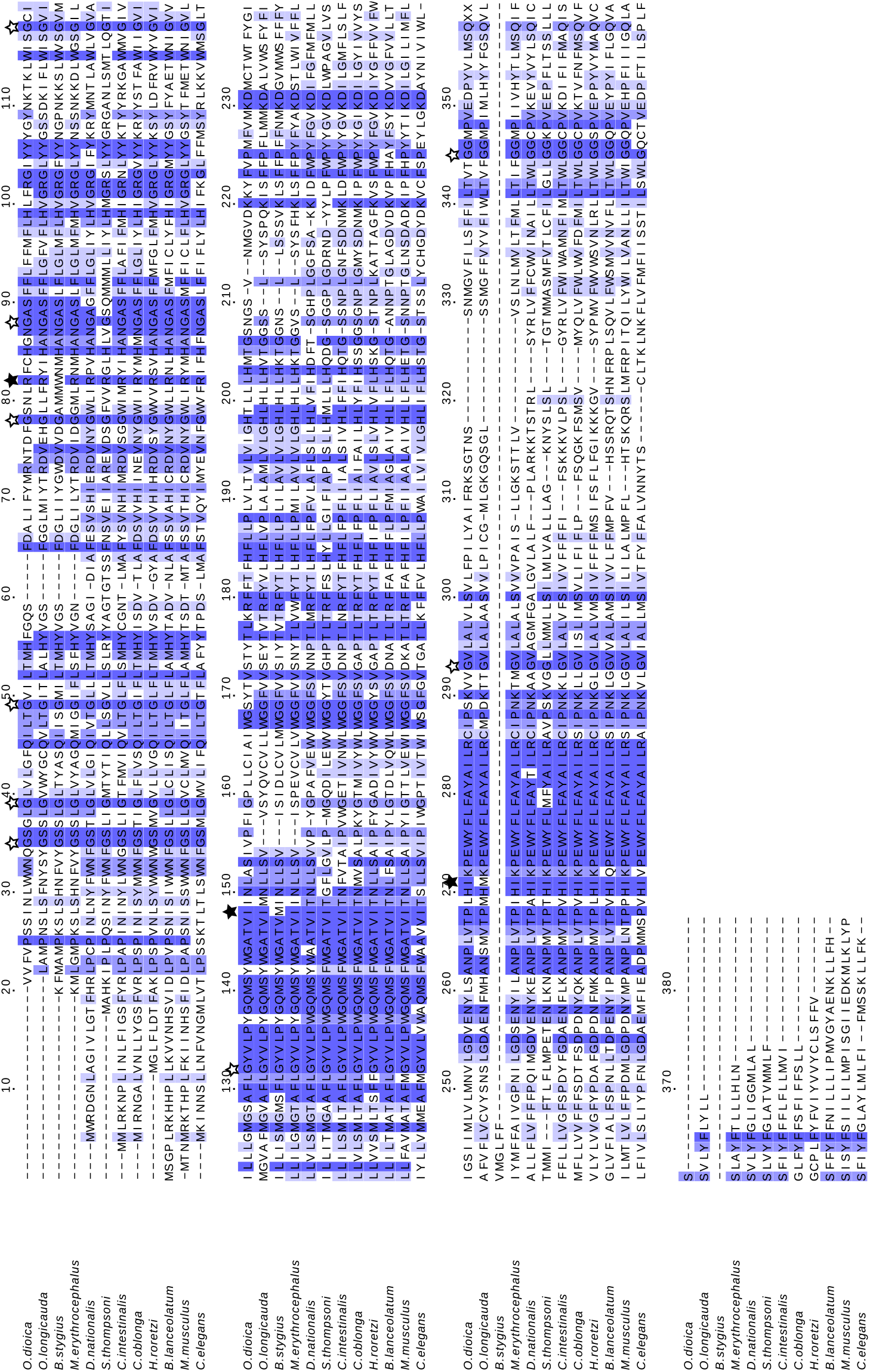
Sequence alignment of Cob proteins. White stars indicate conserved cysteines when at least one tunicate uses an AGR codon. Black stars indicate positions suggesting a different genetic code.

### Possible lineage specific use of ATA Ile and TGA Arg codons

We then searched for positions where a single tunicate species differed from the other sequences with the same replacement amino acid more than once. We found multiple cases of methionine being replaced by isoleucine and arginine replaced by tryptophan in *O. longicauda* and *B. stygius* (Figure 2). Given their phylogenetic proximity, we grouped the two species in the analysis below and we calculated the number of mismatches to the other sequences. We redefined a position as “conserved” if there is at most one mismatch from one sequence to the others.

*M. erythrocephalus* does not seem to use ATA codons and *O. longicauda* and *B. stygius* use ATA codons at positions where all other species had an isoleucine (Ile) (Tables 1 and 2). In the ancestral invertebrate mitochondrial code and the sister vertebrate code, ATA encodes Met. Although Met and Ile both have hydrophobic side chains that often can substitute for each other, this also suggests a change of the genetic code. Evidence for this is that 1) non-appendicularian species do not display ATA codons at positions where all other species encode Ile, 2) the change would be parsimonious as *O. longicauda*, *B. stygius* and *M. erythrocephalus* are more closely related to each other than to *O. dioica* (Naville et al., 2019), and 3) these three species never have ATA codons at positions where Met is conserved in every species (in contrast to *O. dioica*). Furthermore, reversion of the ATA codon to Ile have occurred in other branches of the tree of Life, for instance in echinoderms (Jacobs et al., 1988). Finally, inspection of a partial Cox1 sequence of the related *Bathochordaeus charon* (KT881544.1) provided one extra instance of an ATA codon at a conserved Ile position.

**Table 1:** ATN codons in Cox1 in Oikopleuridae

| lrrrr | *O. dio.* | *O. lon.* | *B. sty.* | *M. ery.* |
| --- | --- | --- | --- | --- |
| ATA number | 22 | 12 | 16 | 0 |
| ATC number | 3 | 8 | 0 | 2 |
| ATG number | 14 | 34 | 25 | 33 |
| ATT number | 29 | 18 | 21 | 38 |
| ATA on cons. Met | 5 | 0 | 0 | 0 |
| ATA on cons. Ile | 0 | 4 | 2 | 0 |

**Table 2:** ATN codons in Cob in Oikopleuridae

| lrrrr | *O. dio.* | *O. lon.* | *B. sty.* | *M. ery.* |
| --- | --- | --- | --- | --- |
| ATA number | 6 | 9 | 9 | 0 |
| ATC number | 5 | 2 | 1 | 2 |
| ATG number | 9 | 9 | 8 | 16 |
| ATT number | 21 | 11 | 11 | 22 |
| ATA on cons. Met | 0 | 0 | 0 | 0 |
| ATA on cons. Ile | 0 | 1 | 1 | 0 |

The TGA codon is known to encode tryptophan (Trp) in vertebrate, invertebrate and ascidian mitochondria (Fox, 1979). We found that *B. stygius* uses TGA at positions where all other species would encode Arg (Tables 3 and 4). This is surprising as these two amino acids are unlikely to functionally substitute for each other. *O. longicauda* does not use TGA codons, and *M. erythrocephalus* does not use TGA at conserved Arg, although it is found at a position where all other species encode for Arg except *C. elegans* which encodes lysine, the other positively charged amino-acid. This again suggests a possible change of genetic code, although the numbers are currently too small to draw a solid conclusion.

**Table 3:** TGR codons in Cox1 in Oikopleuridae

| lrrrr | *O. dio.* | *O. lon.* | *B. sty.* | *M. ery.* |
| --- | --- | --- | --- | --- |
| TGA number | 2 | 0 | 3 | 1 |
| TGG number | 13 | 16 | 19 | 16 |
| TGA on cons. Trp | 1 | 0 | 0 | 0 |
| TGA on cons. Arg | 0 | 0 | 2 | 0 |

**Table 4:** TGR codons in Cob in Oikopleuridae

| lrrrr | *O. dio.* | *O. lon.* | *B. sty.* | *M. ery.* |
| --- | --- | --- | --- | --- |
| TGA number | 3 | 0 | 2 | 1 |
| TGG number | 4 | 7 | 4 | 5 |
| TGA on cons. Trp | 1 | 0 | 0 | 0 |
| TGA on cons. Arg | 0 | 0 | 1 | 0 |

## Discussion

We extracted Cox1 and Cob sequences of four different appendicularians from public databases. As a nucleotide sequence, Cox1 might be useful for mining databases of molecular barcodes sequenced from the environment, or for studies of population diversity within a species. As a protein sequence, it might be useful for refining the phylogeny of appendicularians. However, a translation code needs to be chosen.

Our alignments of tunicate Cox1 and Cob proteins sequences support the view that all tunicates translate AGR codons as Gly (although this conclusion might be limited by the lack of coverage for the Kowalevskiidae and Fritillariidae families). While our analysis suggests that the last common ancestor of the tunicates used the “ascidian” code, it is not possible to conclude that all contemporary tunicates still do, as we found discrepancies on other conserved residues that could be explained by a genetic code change of ATA and TGA codons within a sub-clade of the appendicularians containing *M. erythrocephalus*, *O. longicauda* and *B. stygius*.

The “ascidian” genetic code is table number 13 in the NCBI protein database, where it is used to translate sequences from ascidians and non-ascidian tunicates, for instance *D. nationalis*. However for appendicularians, the NCBI currently applies the invertebrate table (number 5). This has the consequences of turning Gly to Ser at functionally important positions. Therefore, the ascidian is probably a more appropriate default. At present, it is unclear whether some appendicularians have additional changes; however the accurate translation of AGR codons to Gly would nonetheless reduce the amount of error in translated protein sequences.

To confirm a change of genetic code, it is necessary to detect corresponding changes in the respective tRNAs. This beyond reach for the present study because the mitochondrial genomic sequences that we used are extracted from draft genome sequences that may be incomplete, or even contain contaminations (see *B. stygius* in the Methods section). As a result, we also cannot entirely rule out the possibility that we have examined pseudogenes, although the high conservation found in the alignments suggest this in unlikely. For all these reasons, it is necessary to sequence full-length mitochondrial genomes from appendicularians.

## Conclusions

Our alignments of translated mitochondrial sequences suggest that the last common ancestor of living tunicates may have already used the “ascidian” genetic code. Thus, we recommend the use of that code instead of the “invertebrate” one for all tunicates in automatic translation pipelines, with the caveat that additional changes might be found in appendicularians. This observation is a reminder that in biology, exception is the rule, and that each time a mitochondrial sequence is extracted from a species for the first time, it is important to carefully examine its genetic code.

## Supplemental material

Alignment files and description on how we generated them are available in Zenodo at the DOI 10.5281/zenodo.3490310.

## Author contributions

https://dictionary.casrai.org/Contributor_Roles

Conceptualization: JP, CP; Data curation: CP; Formal analysis: JP, CP; Funding acquisition: JP, NML; Investigation: JP, CP; Methodology: CP; Project administration: NML, CP; Resources: n. a.; Software: n. a.; Supervision: NML, CP; Validation: n. a.; Visualization: JP, CP; Writing – original draft: JP, NML, CP; Writing – review & editing: JP, NML, CP.

## Acknowledgements

We thank the OIST’s Scientific Computing & Data Analysis Section for their support, and Ferdinand Marlétaz for critical comments on our manuscript.

